# Prolonged and extended impacts of SARS-CoV-2 on the olfactory neurocircuit

**DOI:** 10.1101/2021.11.04.467274

**Authors:** Megumi Kishimoto-Urata, Shinji Urata, Ryoji Kagoya, Fumiaki Imamura, Shin Nagayama, Rachel A. Reyna, Junki Maruyama, Tatsuya Yamasoba, Kenji Kondo, Sanae Hasegawa-Ishii, Slobodan Paessler

## Abstract

The impact of SARS-CoV-2 on the olfactory pathway was studied over several time points using Syrian golden hamsters. We found an incomplete recovery of the olfactory sensory neurons, prolonged activation of glial cells in the olfactory bulb, and a decrease in the density of dendritic spines within the hippocampus. These data may be useful for elucidating the mechanism underlying long-lasting olfactory dysfunction and cognitive impairment as a post-acute COVID-19 syndrome.

## Results and Discussion

Severe acute respiratory syndrome coronavirus 2 (SARS-CoV-2) has infected over 215 million people, producing an average lethality of 2.1% worldwide^1^. Olfactory dysfunction is one of the first and most common symptoms of the coronavirus disease-2019 (COVID-19)^2^. The proposed mechanism underlying this SARS-CoV-2-induced olfactory dysfunction involves severe damage and impairment of the olfactory epithelium (OE); previous data using animal models indicates apoptosis and desquamation of the entire OE, including olfactory sensory neurons (OSNs)^3^. While the damaged OE is gradually restored in the animal study^4^, many COVID-19 survivors clinically continue to suffer from central nervous system (CNS) symptoms such as depression and memory impairment, as well as chronic olfactory dysfunction in some cases^5^. This study examined the effects of SARS-CoV-2 infection on the CNS using the Syrian golden hamster model, which is a well-established model of COVID-19 that reproduces certain features of human disease^6^.

Olfactory nasal cavity has a complicated structure, divided into multiple regions, e.g. medial/lateral recess on the medial-lateral axis, zone 1-4 areas on the dorsomedial-ventrolateral axis^7^. Each region has a different vulnerability to external harmful agents; dorsomedial side is more vulnerable to methimazole^8^, and lateral side is to lipopolysaccharide^9^. Therefore, we first analyzed the local impact of SARS-CoV-2 on the OE tentatively divided into 4 regions: dorsomedial (DM), dorsolateral (DL), ventromedial (VM), and ventrolateral (VL) region (**Figure 1A**). The hamsters were intranasally inoculated with SARS-CoV-2 at six-weeks of age and samples were collected at several time points. Using immunofluorescence, we detected a significant number of SARS-CoV-2-positive regions throughout the OE at 2 days post-infection (dpi), but not at 8 dpi (**Figure 1A**). Interestingly, SARS-CoV-2-infected cells were observed not only superficially but also deep within portions of the DM region, including the lamina propria. The SARS-CoV-2 antigen was not observed in mature OSNs, as identified by olfactory marker protein (OMP) expression, but in cells around the OSNs, mostly supporting cells (SCs) (**Figure 1B**), which are known to express angiotensin converting enzyme 2 (ACE2), the receptor for SARS-CoV-2^10^. The numbers of SCs significantly decreased in the VM and VL and could not be determined in the DM due to the complete loss of the OE at 5 dpi, although the damage was recovered almost completely in all regions by 21 dpi (**Figure 1C**). Interestingly, no SARS-CoV-2 antigen was detected within the sagittal section of the whole brain (**Figure 1-figure supplement 1A**), including the olfactory bulb (OB, **Figure 1-figure supplement 1B**) and hippocampus (**Figure 1-figure supplement 1C**). The data from our study suggest that the SARS-CoV-2 did not infect the brain parenchyma or that the level of infection was below detection limit, however, previous research revealed that SARS-CoV-2 RNA and viral antigen can be detected in the brain^11^. Moreover, we found that the OE thickness transiently decreased at 5 dpi but recovered fully by 21 dpi, as was the case for the SC numbers (**Figure1, Figure 2-figure supplement 1A,B**). Nevertheless, the density of mature OSNs did not completely recover up to 42 dpi, suggesting that the maturation of OSNs may be delayed and/or incomplete (**Figure 2-figure supplement 1A,C**).

**Figure 1.**
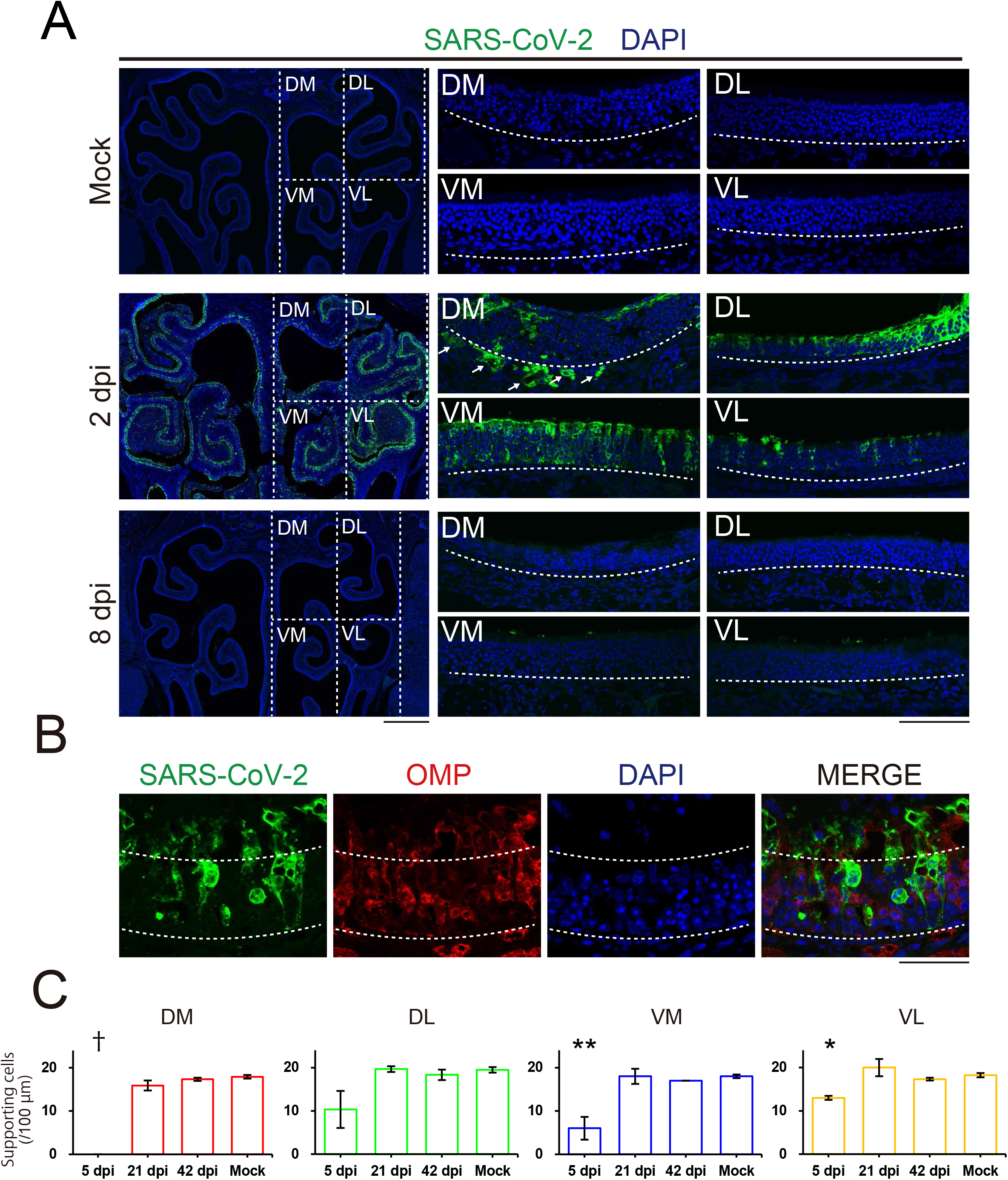
Distribution of severe acute respiratory syndrome coronavirus 2 (SARS-CoV-2) in the OE. A, Distribution of SARS-CoV-2 in the olfactory epithelium (OE) at mock, 2 dpi, and 8 dpi. The OE was divided into 4 regions: dorsomedial (DM), dorsolateral (DL), ventromedial (VM), and ventrolateral (VL). SARS-CoV-2-positive cells were localized in the 4 regions of the OE at 2 dpi. The white arrows indicate the SARS-CoV-2-positive cells in the lamina propria. Scale bars, 1 mm (left), 100 μm (right). B, Profile of SARS-CoV-2 infection in the OE. The apical white dashed line indicates the surface of the epithelium. The basal white dashed line is the border of the OE and lamina propria. Scale bar, 50 μm. C, Numbers of the supporting cells (SCs) in the OE in DM, DL, VM, and VL. (one-way ANOVA followed by Dunnett’s post hoc test. *p < 0.05, **p < 0.01, ***p < 0.001, † means unevaluable as the OE was completely desquamated and SCs were not detectable). Number of samples; n = 3 (5, 21 and 42 dpi) and 4 (mock).

Our data indicate that the DM is the most vulnerable to SARS-CoV-2 in the OE. This is reasonable, as ACE2 receptors are predominantly expressed in this region^12^. The DM is considered to be included in the zone 1 of the zonal categorization^7^. Interestingly, the OSNs in zone 1 only express the endocellular enzyme NAD(P)H quinone oxido-reductase 1 (NQO1)^13^. NQO1 acts as an antioxidant but also mediates ROS generation, being involved in increasing certain forms of neuronal damage after injury^8^. This led us to analyze the relationship between NQO1 expression patterns and neural damage caused by SARS-CoV-2 infection. The numbers of OMP-positive cells decreased after infection and were not completely restored in both NQO1-positive and -negative regions (**Figure 2A,B,C**). However, the damage was more severe in NQO1-positive regions than in NQO1-negative regions (**Figure 2D**). Furthermore, the presence of macrophages, which are labeled using the ionized calcium binding adaptor molecule 1 (Iba1) suggest prolonged activation in the lamina propria within NQO1-positive regions (**Figure 2E,F,G**). In contrast, macrophages quickly returned to a normal level within NQO1-negative regions. These data suggest that NQO1-positive areas were more susceptible to SARS-CoV-2 than the negative areas. These trends were also observed in the next olfactory relay center, the OB, in which the OSNs have synaptic contact with multiple neurons within the glomerulus (**Figure 3A,B**).

**Figure 2.**
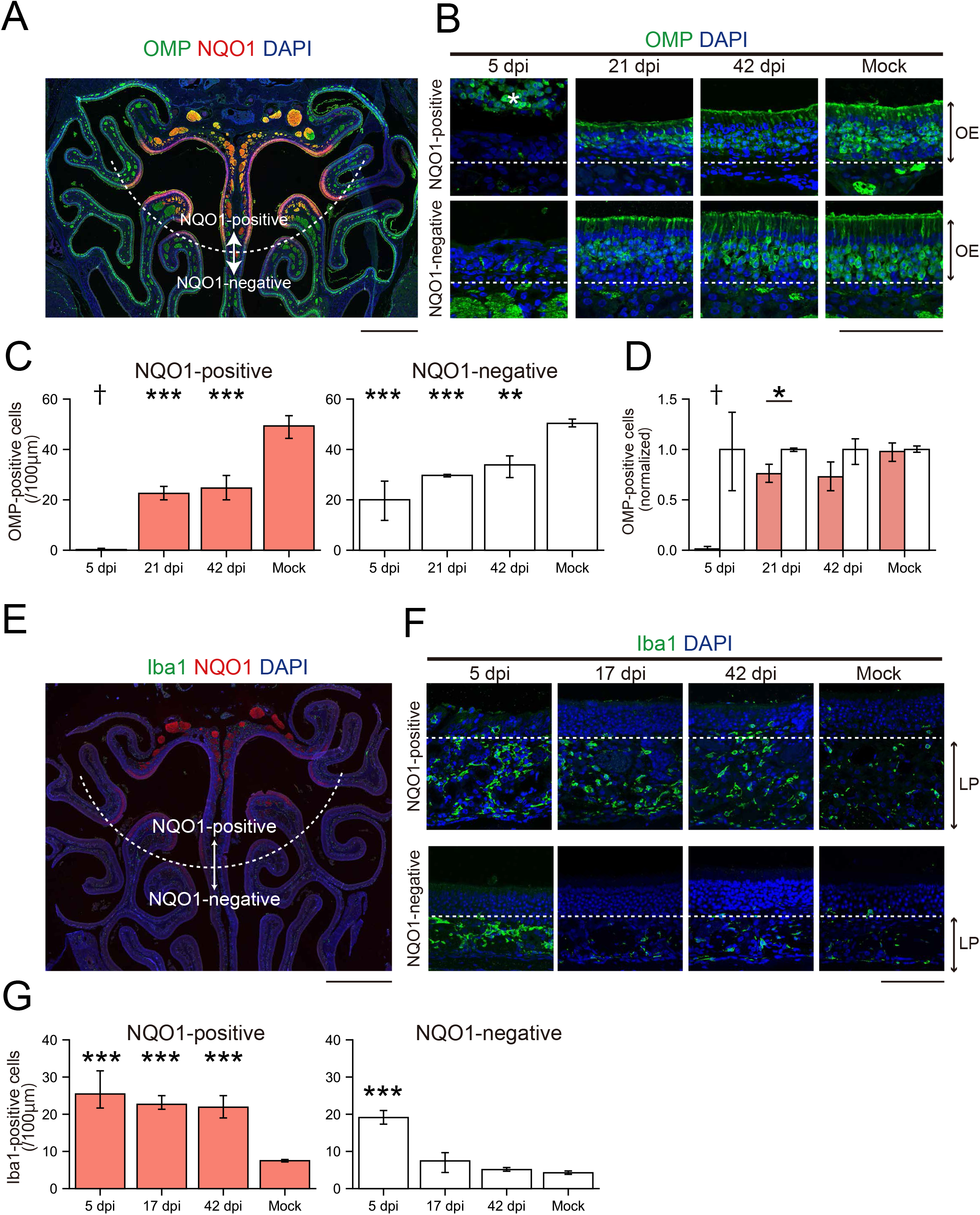
Region-specific damage of the olfactory epithelium (OE) due to SARS-CoV-2 infection. A, Representative coronal section of the OE stained with olfactory marker protein (OMP), NAD(P)H quinone oxido-reductase 1 (NQO1), and DAPI. The white dashed line is the border of the NQO1-positive or -negative region. Scale bar, 1 mm. B, Representative images of OMP-positive cells at 5, 21, and 42 dpi or mock in the OE. The white dashed line is the border between the OE and lamina propria. The asterisk indicates desquamated epithelium. Scale bar, 100 μm. C, Numbers of OMP-positive cells in the NQO1-positive or -negative region. (one-way ANOVA followed by Dunnett’s post hoc test. **p < 0.01, ***p < 0.001, † means unevaluable as the OE was completely desquamated and OMP-positive cells were not countable). Number of samples: n = 3 (5, 21 and 42 dpi) and 4 (mock). D, Comparison of the normalized density of OMP-positive cells in NQO1-positive and NQO1-negative regions. (n = 3, Welch’s t-test. *p < 0.05, † means unevaluable as the OE was completely desquamated and OMP-positive cells were not countable). E, Representative coronal section of the OE stained with ionized calcium binding adaptor molecule 1 (Iba1), NQO1, and DAPI. The white dashed line is the border of the NQO1-positive or -negative region. Scale bar, 1 mm. F, Representative images of Iba1-positive cells at 5, 17, and 42 dpi or mock. The white dashed line is the border between the OE and lamina propria. Lamina propria, LP. Scale bar, 100 μm. G, Densities of Iba1-positive cells in the NQO1-positive or -negative region. (one-way ANOVA followed by Dunnett’s post hoc test. ***p < 0.001). Number of samples: n = 3 (5, 17 and 42 dpi) and 4 (mock).

**Figure 3.**
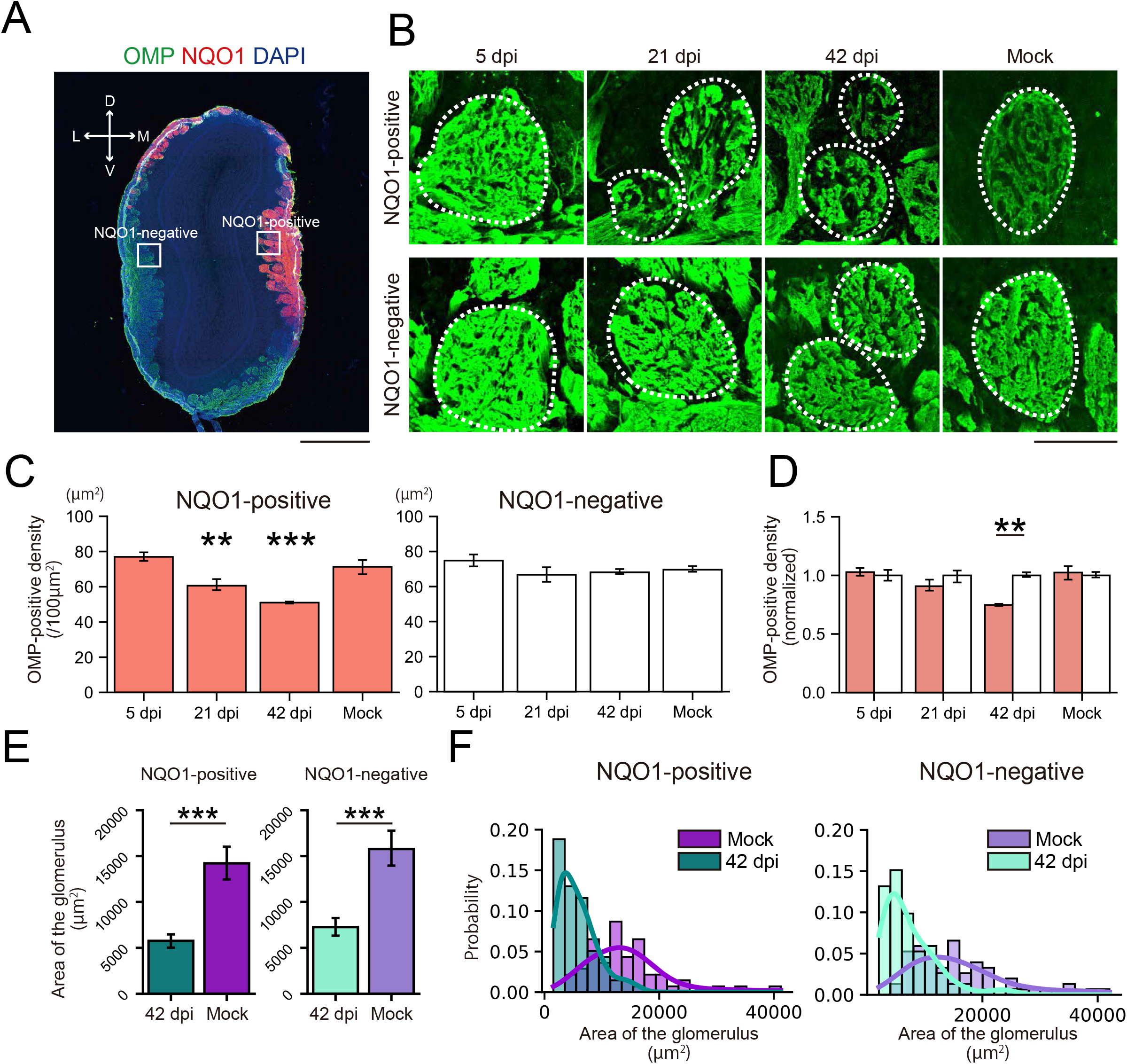
Region-specific changes in the glomeruli in the olfactory bulb (OB) due to SARS-CoV-2 infection. A, Representative coronal section of the OB stained with olfactory marker protein (OMP), NQO1, and DAPI. Scale bar, 1 mm. B, Representative time series of glomeruli stained with OMP. Each circled area corresponds to a glomerulus. Scale bar, 100 μm. C, The density of OMP-positive areas within the glomerulus compared between NQO1-positive or - negative regions. (n =3, one-way ANOVA followed by Dunnett’s post hoc test. **p < 0.01, ***p < 0.001). D, Comparison of the normalized OMP-positive area in NQO1-positive and NQO1-negative regions. (n = 3, Welch’s t-test. **p < 0.01). E, The size of the glomerulus at 42 dpi and mock. (Welch’s t-test. ***p < 0.001). Number of samples; n = 78, 60, 86, and 67 glomeruli from 42 dpi of NQO1-positive, mock of NQO1-positive, 42 dpi of NQO1-negative, and mock of NQO1-negative, n = 3 animals in individual group. F, Histogram of the area of the glomerulus. The regression curves were fitted using kernel density estimation.

In the OB, the density of the OSN axon terminal of each glomerulus was significantly decreased within NQO1-positive regions, but not in the NQO1-negative regions (**Figure 3C,D**). Interestingly, the size of the glomeruli themselves decreased not only in NQO1-positive but also in NQO1-negative regions (**Figure 3E,F**). These data indicate that SARS-CoV-2 infection impacts odor information processing within the whole OB, but especially prominent in NQO1-positive regions. Next, we carefully examined the effects of SARS-CoV-2 on the profile of glial activities (**Figure 4A, Figure 4-figure supplemental 1A, Figure 5-figure supplement 1A**) throughout the multiple layers of the OB: the olfactory nerve layer (ONL) where the OSN axon shaft are densely packed, the glomerular layer (GL) where the OSN axon synapse to the relay neurons, and the external plexiform layer (EPL) where the projection neurons interact with the local interneurons. Remarkably, the Iba1 signal peaked at 17 dpi (**Figure 4B**), in contrast to the OE, in which it peaked at 5 dpi, indicating that microglia in the OB became activated more slowly than macrophages in the OE. Interestingly, the ONL, occupied by OSN axons, showed significantly strong microglial activation, even at 5 dpi. This indicates that the microglia in the ONL respond to the damage of the OSN axons from the OE quickly and directly, and this damage would provide an indirect secondary impact on the postsynaptic circuits in the GL and EPL. Consistent with the impact of SARS-CoV-2 on the OSN axon terminals within the OB (**Figure 3**), the impact on the NQO1-positive regions was significantly stronger within the ONL at 5 dpi and within the GL at 17 dpi, as compared to that of the NQO-1-negative regions (**Figure 4B,D,F**). However, the area occupied by glial fibrillary acidic protein (GFAP)-expressing reactivated astrocytes was enlarged at 17 dpi within the GL (**Figure 4-figure supplemental 1B,C,E**), although the dynamics of GFAP-positive cells were not different based on NQO1 expression (**Figure 4-figure supplemental 1D,F**). This indicates that astrocyte activity is strongly associated with the activity of microglia in the GL compared to that in other layers.

**Figure 4.**
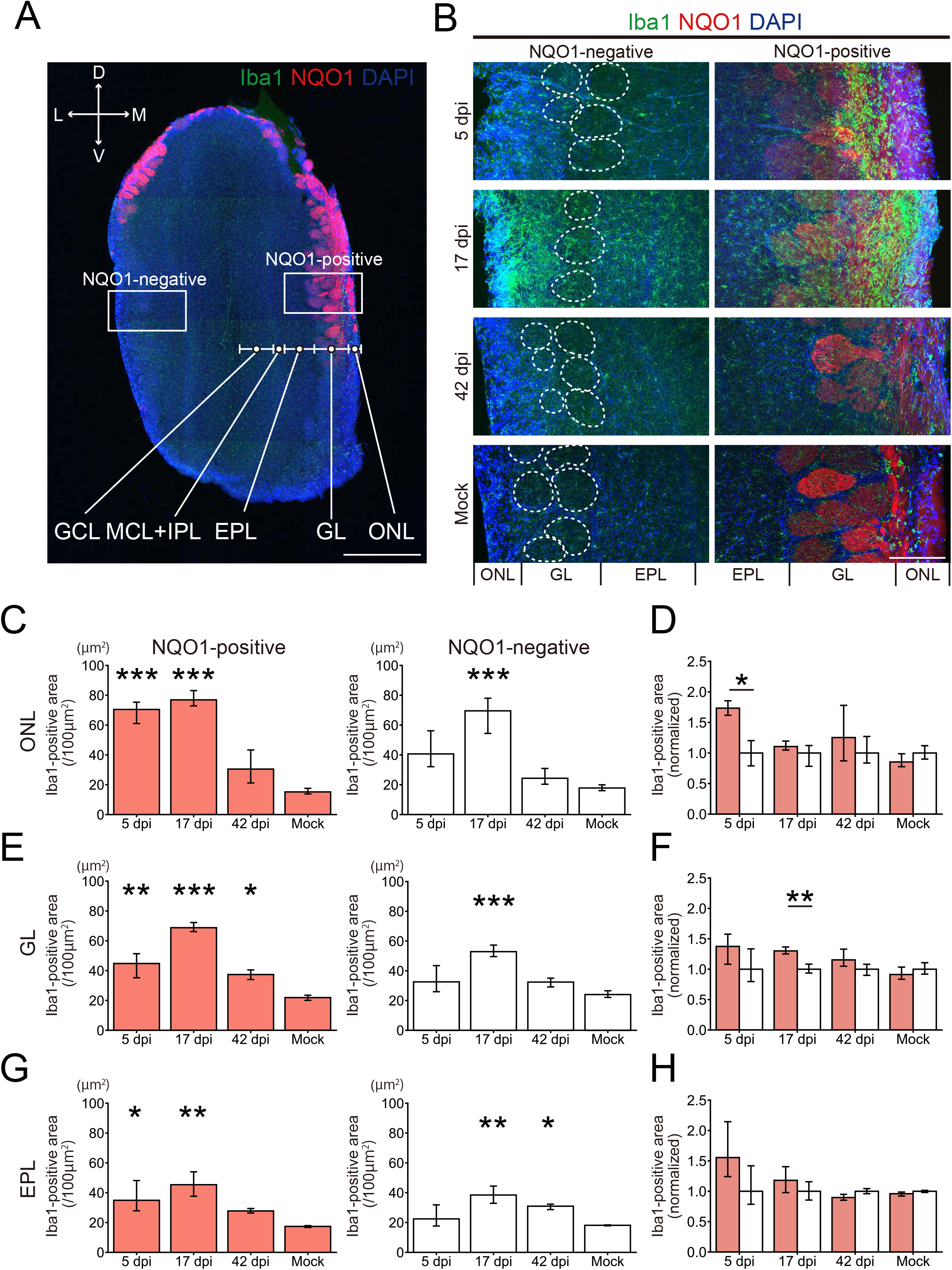
Region-specific microglial activation in the OB due to SARS-CoV-2 infection. A, Representative coronal section of the OB stained with Iba1, NQO1, and DAPI. The OB has a multi-layered structure from surface to the center; Olfactory nerve layer (ONL), Glomerular layer (GL), External plexiform layer (EPL), Mitral cell layer (MCL), Internal plexiform layer (IPL), and Granule cell layer (GNL). Squares are representative regions in the analysis of NQO1-positive and -negative region. Scale bar, 1 mm. B, Representative images of ONL, GL, and EPL stained with Iba1, NQO1, and DAPI. White dashed circles indicate glomeruli. Scale bar, 200 μm. C,E,G, Iba-1-positive area in the ONL (c), GL (e), and EPL (g). (n = 3, one-way ANOVA followed by Dunnett’s post hoc test. *p < 0.05, **p < 0.01, ***p < 0.001). D,F,H, Comparison of the normalized values of the Iba1-positive area in NQO1-positive and NQO1-negative regions in the ONL (d), GL (f), and EPL (h). (n = 3, Welch’s t-test. *p < 0.05, **p < 0.01).

Next, we examined the impact of SARS-CoV-2 on higher brain areas, including the piriform cortex (PC) and the hippocampus (**Figure 5, Figure 5-figure supplemental 1**). In the PC, which is one of the olfactory cortices receiving direct inputs from the OB, GFAP-expression appeared transient in the pia mater but persisted in the layer 1 of the PC up to 42 dpi (**Figure 5-figure supplemental 1B**). Activated microglia/macrophages cells labeled by Iba1 within the pia mater increased in numbers at 5 and 17 dpi, although the Iba1-positive area was unchanged in layer 1 (**Figure 5-figure supplemental 1C,D**). The peak of this activity was not correlated to the presence of SARS-CoV-2 within the nasal cavity (**Figure 1A**), indicating that glial activation in the PC may not be induced directly by the virus through the olfactory pathway but by local and/or systemic inflammatory responses. In fact, inflammatory cytokines are systemically elevated in both severe and non-lethal COVID-19 models^14^. Given that ACE2 is expressed in the Bowman’s glands of the lamina propria^15^ and that macrophages there were activated after SARS-CoV-2 infection (**Figure 2F,G**), the immune response in the lamina propria may induce the elevation of intravascular inflammatory cytokines, triggering the activation of glial cells and macrophages in the CNS. In the hippocampus, GFAP-positive astrocytic endfeet were easily detectable around the blood vessels of the apical dendritic region at 5, 8, 17, and 42 dpi (**Figure 5B**), as compared to those of the basal region. The peak Iba1-signal in the basal region was achieved by 8 dpi, with the area being restored by 42 dpi (**Figure 5C,D**). In contrast, the Iba1-signal in the apical region remained strong until 42 dpi (**Figure 5C,D,E**). These data indicate that SARS-CoV-2 infection in the nostril triggered the activation of microglia and astrocytes, even in the hippocampus, and that the impacts are significantly different in individual layer. Thus, an interesting question could be posed as to whether the activated microglia and astrocytes could induce any changes in the neuronal circuits?

**Figure 5.**
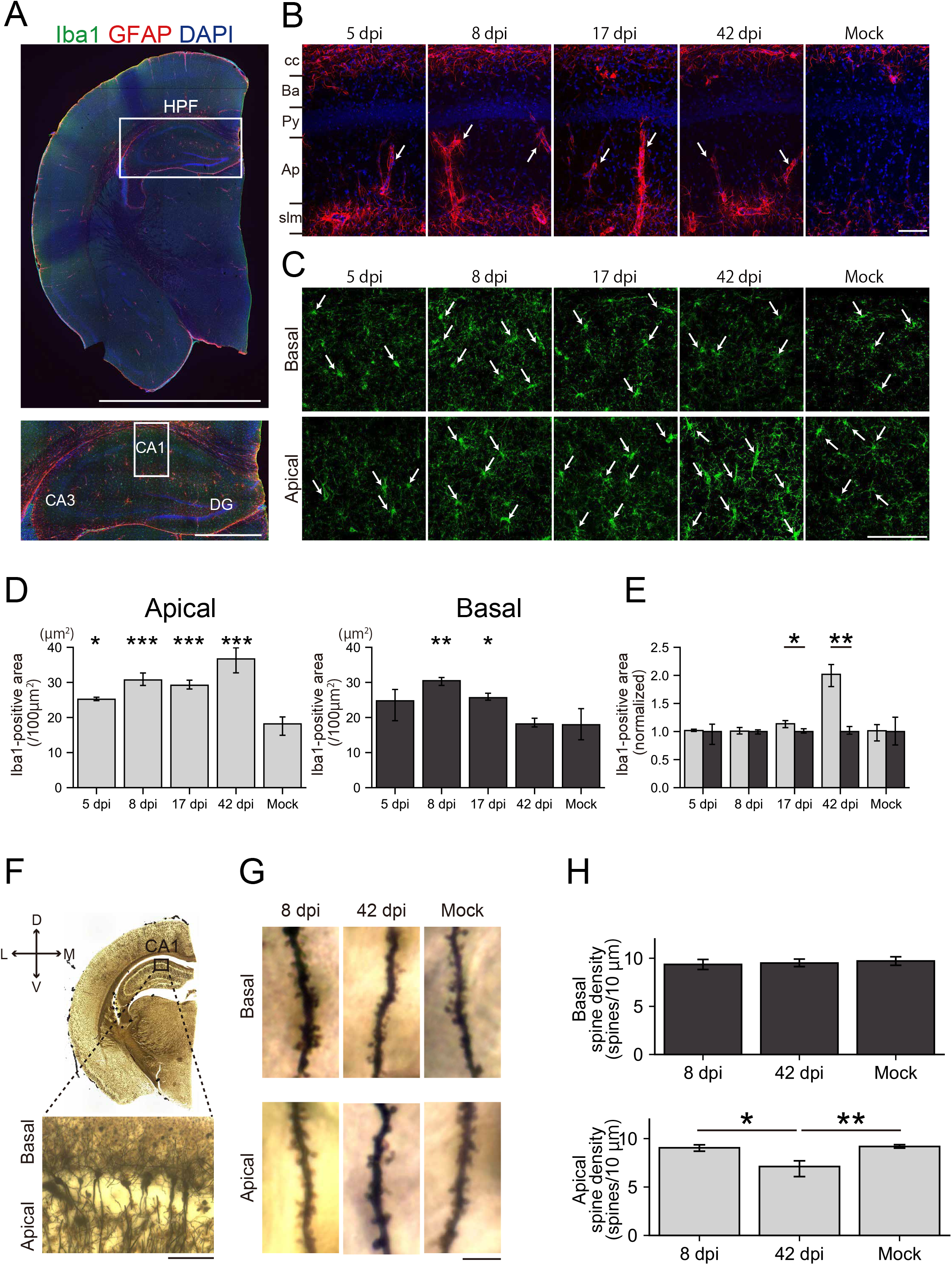
Glial activation and loss of dendritic spines in the hippocampus due to SARS-CoV-2 infection. A, The upper image is a representative coronal image of the cerebral hemisphere stained with Iba1, GFAP, and DAPI. The upper white square indicates the hippocampal formation (HPF). The lower panel is a high-magnification image. The lower white square-labeled region (CA1) is enlarged in Figure 5B. Scale bar, 5 mm (upper), 1 mm (lower). B, Representative image of astrocytes in the CA1 stained with glial fibrillary acidic protein (GFAP) and DAPI at 5, 8, 17, 42 dpi, and mock. Arrows indicate GFAP-positive astrocytic endfeet. Corpus callosum, cc; Basal dendrite layer, Ba; Pyramidal cell layer, Py; Apical dendrite layer, Ap; stratum lacunosum-moleculare, slm. Scale bar, 100 μm C, Representative image of microglia in the basal and apical region of the CA1 stained with Iba1. Arrows indicate Iba1-positive microglia. Scale bar, 100 μm D, Area of Iba1-positive cells in the apical and basal regions of the CA1. (n = 3, one-way ANOVA followed by Dunnett’s post hoc test. *p < 0.05, **p < 0.01, ***p < 0.001). E, Comparison of the normalized density of Iba1-positive cells in the NQO1-positive and NQO1-negative regions. (n = 3, Welch’s t-test. *p < 0.05, **p < 0.01). F, Upper panel is a representative image depicting Golgi-impregnated neurons of the cerebral hemisphere. Lower panel is a high-magnification image of the CA1. Scale bar, 100 μm. G, Representative image depicting Golgi-impregnated dendrites and dendritic spines in the apical and basal region of the CA1. Scale bar, 5 μm. H, Densities of apical and basal dendritic spines. (n = 3, one-way ANOVA followed by Tukey’s post hoc test. *p < 0.05, **p < 0.01, neurons = 24, 48, 46; spines = 1610, 1789, 2453 from the basal dendrites of 8, 42 dpi, and mock group, neurons = 25, 41, 43; spines = 1252, 1262, 1317 from the apical dendrites of 8, 42 dpi and mock group).

There are many reports that reveal glial cells induce synaptic modulation^16^, synaptic loss^9^, synaptic plasticity^17^, and change of synaptic density^18^. These changes may be associated with dementia^19^. Therefore, we analyzed dynamics of spine density within the hippocampus using Golgi stained brain sections (**Figure 5F,G**). Basal dendritic spines in the hippocampal CA1 were extremely stable. In contrast, the density of apical dendritic spines was significantly decreased at 42 dpi (**Figure 5H**) This finding may be associated with the prolonged activation of microglial cells, most notably significant at 42 dpi (**Figure 5E**). These results suggest that intranasal inoculation of SARS-CoV-2 induces glial cell activation and changes spine density within the higher brain regions, including the hippocampus. Future studies should determine whether these changes impact animal behavior and if so, how long it is required for recovery.

In summary, we report that single priming with SARS-CoV-2 resulted in long-lasting effects in hamsters, not only on the OE but also within the brain regions critical for cognitive function. We found drastic CNS changes including glial activation and synaptic dynamics, without any SARS-CoV-2 antigen by immunostaining. Our study may be of importance for better understanding the mechanism(s) driving the olfactory impairment and potentially cognitive dysfunction present in COVID-19 survivors.

## Methods

### Cell and virus

Vero E6 cells were maintained using Dulbecco’s modified Eagle’s medium (DMEM) supplemented with 10% fetal bovine serum (FBS), 1% penicillin-streptomycin, and L-glutamine. SARS-CoV-2 (USA/WA-1/2020) was propagated in Vero E6 cells with DMEM supplemented with 2% FBS. Cell culture supernatant was stored in the −80°C freezer until use.

### Animal experiments

Six-week-old female Syrian golden hamsters were purchased from Charles River. Syrian golden hamsters were anesthetized with isoflurane and inoculated bilateral-intranasally with 10^5^ 50% tissue culture infectious dose (TCID50) of SARS-CoV-2 diluted in 100 μl of phosphate-buffered saline (PBS) or PBS as a mock control. Mock controls were sacrificed 42 days after inoculation. All hamsters were housed in the animal biosafety level-2 (ABSL-2) and ABSL-3 facilities within the Galveston National Laboratory at the University of Texas Medical Branch (UTMB). All animal studies are reviewed and approved by the Institutional Animal Care and Use Committee at UTMB and are conducted according to the National Institutes of Health guidelines.

### Immunohistochemistry

Hamsters were euthanized using a high-flow rate of CO_2_ followed by thoracotomy at each time point to collect samples. The heads of hamsters were fixed in 10% buffered formalin for 7 days before removal from the BSL-3 facility and followed by incubation in phosphate buffered saline (PBS). After OBs were extracted from the skull, right OBs were sectioned into 100 μm thick sections using a vibratome and left OBs were proceeded to the tissue Golgi-Cox procedures. The nasal tissues including the olfactory epithelium were decalcified with 10% EDTA (pH 7.0) for 7 days at 37 °C with gentle shaking. Decalcification samples were dehydrated in 100% ethanol three times and embedded in paraffin. Serial coronal sections (5 μm thick) were cut and mounted on glass slides. Deparaffinized sections and free-floating brain slices were incubated for 10-30 min in Target Retrieval Solution (S1700; Dako) at 72-102 °C in a water-bath for antigen retrieval^20^. Non-specific antibody binding was blocked by Serum-free protein block (Dako). Samples were incubated for 12 hr in a solution containing the following primary antibodies: olfactory marker protein (OMP, goat polyclonal, 1:5000 dilution; Wako Chemicals), SARS-CoV-2 Nucleocapsid antibody (rabbit polyclonal, 1:100; Sino Biological), anti-Iba1 (rabbit polyclonal, 1:300; Wako), anti-Iba1 for paraffin section (rabbit polyclonal, 1:300; Wako), anti-Iba1 (goat polyclonal, 1:300; Abcam), anti-NQO1 (rabbit polyclonal, 1:300; Cell Signaling), anti-NQO1 (goat polyclonal, 1:300; Abcam), and anti-GFAP antibody (mouse monoclonal, 1:500; Millipore). Unbound antibodies were removed by washing for 3 min with PBS or 30 min with PBS containing 0.1% Triton X-100, respectively. Primary antibodies were detected by incubation for 1 hr with a solution containing the following secondary antibodies; goat anti-rabbit Fluor 488, goat anti-mouse Fluor 568, donkey anti-goat Alexa Fluor 488, and donkey anti-rabbit Alexa Fluor 568 (1:200; Invitrogen). To visualize cell nuclei, DAPI or Hoechst 33342 (1:500; Thermo Fisher) was applied during the secondary antibody stain. Samples were mounted in Vectashield hard-set mounting medium (Vector Laboratories) or 70% Glycerol.

### Golgi-Cox staining

To quantify the number of dendritic spines, brain slices were treated with the Rapid GolgiStain Kit according to the manufacturer’s instructions (ND Neurotechnologies, Inc., PK401) with minor modifications. Since sample conditions, especially after formalin fixation, were significant and critical variables for Golgi staining^21–24^, modified Golgi-Cox staining was performed as described in previous reports^24,25^. Briefly, the brain slices were incubated in the impregnation solution for 14 days in 37°C.

The impregnation solution was mixed using a 1:1 ratio of Solutions A and B, which contain mercuric chloride, potassium dichromate, and potassium chromate. The solution was prepared 24 hr before use, to allow precipitate formation, and covered with aluminum foil and kept in 37°C. The sample was replaced in a sucrose-based solution (solution C) for 10 min at room temperature. The sections were transferred and agglutinated onto gelatin-coated slides. The sections were rinsed twice for 5 min each in distilled water to remove the impregnation solution. The sections were then incubated in the staining solution (Solution D: Solution E: distilled water, 1:1:2) for 10 min at room temperature.

### Analysis of immunohistochemistry

To minimize the anatomical variations of the nose, three coronal sections covered with olfactory neuroepithelium were analyzed every 500 μm intervals. The OE consists of three cell types: olfactory sensory neurons, supporting cells, and basal stem cells. For each OE, coronal sections of the OE were divided into 4 regions: dorsomedial (DM), dorsolateral (DL), ventromedial (VM) and ventrolateral (VL). The thickness of olfactory epithelium was measured between the surface of neuroepithelium and the line of basal cells. We counted the numbers of OMP-positive cells for mature olfactory sensory neurons. To evaluate the numbers of supporting cells (SCs), we defined and counted the SCs as the columnar as the cells located proximal to the nasal cavity, as indicated in a previous report^26^. To investigate the presence of macrophages in the lamina propria, Iba1-positive cells were counted. All analyses were performed for each group per 100 μm OE length. Three regions were randomly selected, with the regions being at least 100 μm apart, and the averaged values were used. To quantify the densities of the Iba1- and GFAP-positive area, we defined the positive area as one that exceeds two SDs of the mean background intensity. Subsequently, binary images were overlapped onto the original images and the densities were automatically measured using ImageJ software. OMP-positive cells (**Figure 2D**), Iba1-postive areas (**Figure 4D,F,H**), OMP-positive areas (**Figure 3D**), and GFAP-positive areas (**Figure 4-figure supplement 1D,F**) were adjusted to normalize the average values in the NQO1-negative. Iba1-positive areas in the hippocampus (**Figure 5E**) were adjusted to normalize the values in the basal region.

### Analysis of Dendritic Spine Density

The location of CA1 was identified using a previously published brain map^27^. Pyramidal neurons in the CA1 were selected and three apical and three basal dendritic segments were analyzed^28^. For analysis of Golgi-impregnated neurons, the criteria are the following, according to a previous report^29^: (1) dendrites with a length of at least 50 μm, (2) consistent and dark impregnation along the entire extent of dendrites, and (3) relative isolation from neighboring impregnated dendrites. Densities of dendritic spines were calculated as the mean numbers of spines per 10μm per dendrite per neuron in individual animals per group.

### Microscopy

Images (**Figure 5F,G, Figure 1-figure supplement 1A, and Figure 2-figure supplement 1A**) were captured using a digital CCD (Keyence BZ-X800, Japan) with 20 × (NA = 0.45), 100 × (NA = 1.45) objective lenses. The size of 20 × single horizontal images (**Figure 5F, Figure 1-figure supplement 1A, and Figure 2-figure supplement 1A**) were set to 1920 × 1440, with pixel size of 0.37 × 0.37 and z-spacing of 5 μm. The size of 100 × single horizontal images (**Figure 5G**) were set to 1920 × 1440, with pixel size of 0.07 × 0.07 and z-spacing of 0.2 μm. For samples labeled with multiple antibodies (**Figure 1A,B, Figure 2A,B,E,F, Figure 3A,B, Figure 4A,B, Figure 5A,B,C, Figure 1-figure supplement 1B,C, Figure 2-figure supplement 1A, Figure 4-figure supplement 1A,B, and Figure 5-figure supplement 1A,B**), images were obtained using a confocal microscope (Zeiss LSM 880, Germany) with a 20 objective lens (NA = 0.8). Fluorophores were excited by 405, 488, 561 nm lines of diode lasers. The sizes of single horizontal images were set to 512 × 512, with pixel size of 0.83 × 0.83 and z-spacing of 1 μm.

### Statistical analysis

To prevent arbitrary analysis, all data were randomly and blindly analyzed by two trained human operators. R (https://www.r-project.org/) and custom-written Python scripts were used for statistical analysis. Mean ± SD was used to indicate biological variations. In Figure2D, Figure 3D,E, Figure 4D, F, H, Figure 5E, Figure 4-figure supplemental 1D, and Figure 4-figure supplemental 1F, Welch’s t-tests were used. Compared to mock groups, comparisons among multiple groups (Figure 1C, Figure 2C,G, Figure 3C,E, Figure 5D, Figure 2-figure supplement B,C, Figure 4-figure supplemental C,E, and Figure 5-figure supplement 1C,D) were evaluated with one-way ANOVA and followed by Dunnett’s post hoc test. In Figure 5H, one-way ANOVA and followed by Tukey’s post hoc test were performed. P-values for figures are depicted as follows: *p < 0.05, **p < 0.01, ***p < 0.001.

## Acknowledgements

We would like to thank both the histology core for the HE-staining and imaging core for use of the microscope at UTMB, C.A. Grant, M.A. Micci, K. Suenaga, and H. Iigusa for technical support in preparation of the manuscript, as well as R. Kamei T.Higashi and Y. Kashiwagi for discussions. We also thank T. Makishima for critical reading of the manuscript. This project was funded by John S. Dunn Endowment to S.P.

## Author contributions

J.M. designed the study. R.A.R. and J.M. collected the samples. M.K.U. and S.U. designed the experiments and performed the experiments and imaging. M.K.U, S.U., R.K., and K.K. quantified and M.K.U, S.U., and S.H.I. analyzed data. M.K.U, S.U., R.K., K.K., S.H.I, F.I., S.N., T.Y., and S.P. interpreted data. S.U. designed the statistical analysis. S.H.I., F.I., S.N., R.K., T.Y., K.K. and S.P supervised the experiments. M.K.U. and S.U. co-wrote the initial manuscript. S.H.I., F.I., S.N., R.K., K.K., J.M., R.A.R., T.M., T.Y. and S.P. revised the manuscript. S.P. conceived the study, designed experiments, interpreted data, and co-wrote the manuscript with input from all authors.

## Figure supplements legends

**Figure 1-figure supplement 1.**
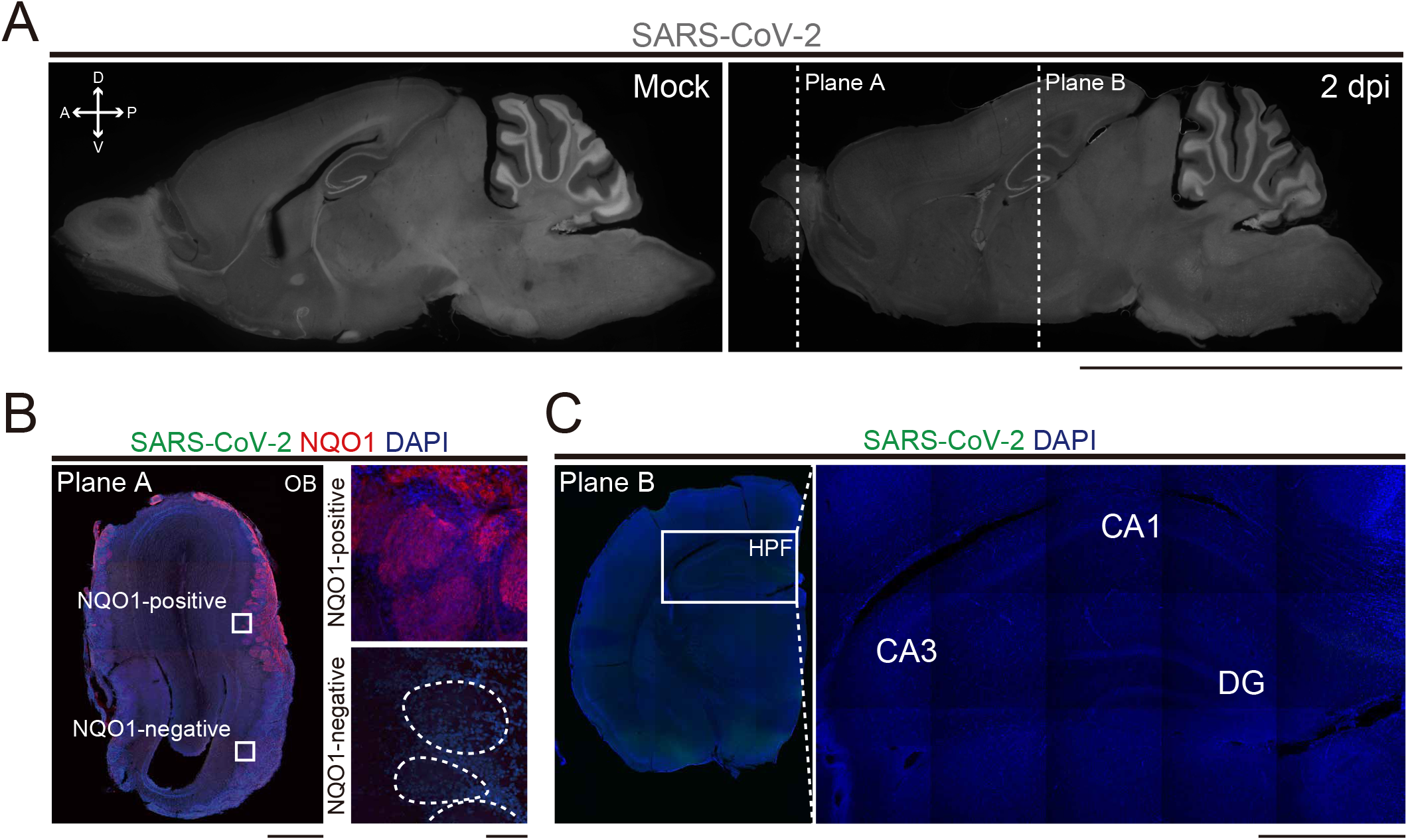
Distribution of SARS-CoV-2 in the OB. A, Sagittal sections of the whole brain stained by SARS-CoV-2 in mock and 2 dpi. Planes A and B are shown as coronal sections in the Figure B and C. Scale bar, 1 cm. B, Representative image of the coronal section of the OB. Right panels are high-magnification images of the GL in NAD(P)H quinone oxido-reductase 1 (NQO1)-positive (upper) and -negative (lower) regions. White dashed circles indicate glomeruli. Scale bars, 1 mm (left), 100 μm (right). C, Left panel shows a representative image of the coronal section of the cerebral hemisphere. The white box indicates hippocampal formation (HPF). Right panel is a high-magnification image of the HPF; CA1, CA3, and DG. Scale bar, 1 mm.

**Figure 2-figure supplement 1.**
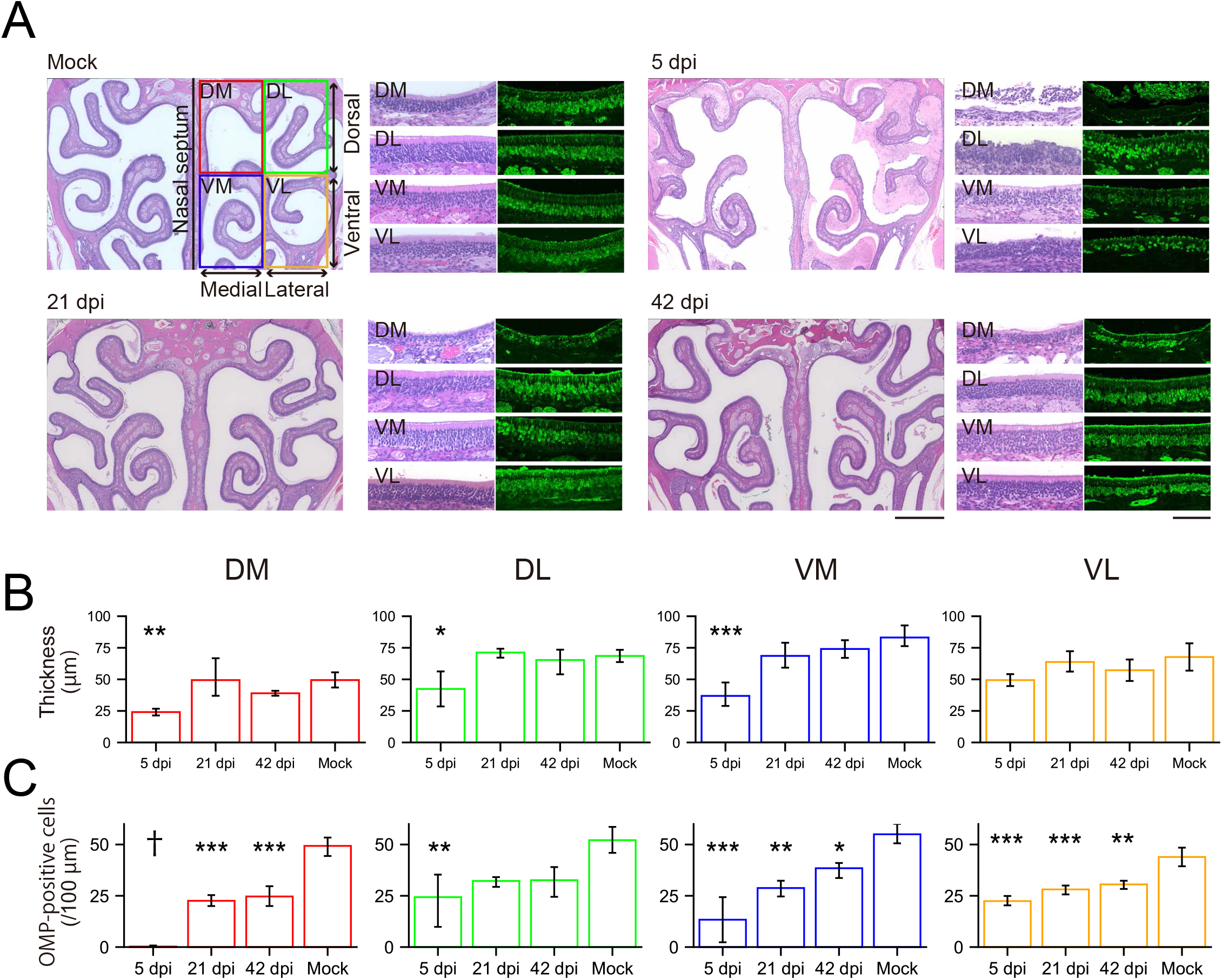
Cellular properties in the OE due to SARS-CoV-2 infection. A, Representative images of the OE stained with HE and OMP at 5 dpi, 21 dpi, and 42 dpi and mock,. Scale bars, 1 mm (left), 100 μm (right). B, The thickness of the OE in 4 subdivided areas at 5, 21, and 42 dpi and mock; DM, DL, VM, and DL. (one-way ANOVA followed by Dunnett’s post hoc test. *p < 0.05, **p < 0.01, ***p < 0.001). Number of samples; n = 3 (21 dpi) and 4 (5, 42 dpi, and mock). C, Numbers of the OMP-positive cells in the neuroepithelium at 5, 21, and 42 dpi and mock; DM, DL, VM, and DL. (n = 3, one-way ANOVA followed by Dunnett’s post hoc test. *p < 0.05, **p < 0.01, ***p < 0.001, † means unevaluable as the OE was completely desquamated and OMP-positive cells were not countable).

**Figure 4-figure supplement 1.**
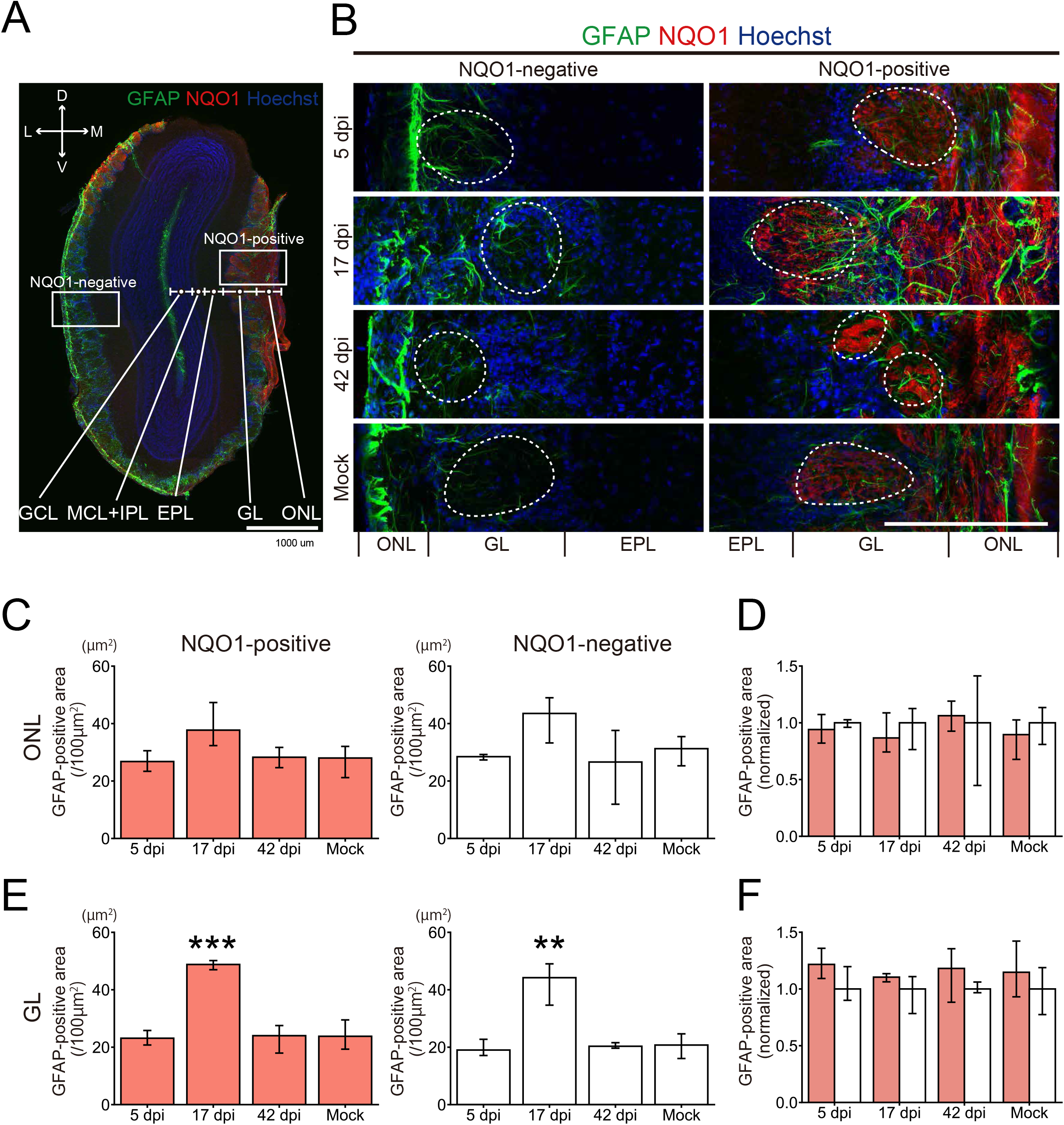
GFAP-positive cells in the OB. A, Representative coronal section of the OB stained with glial fibrillary acidic protein (GFAP), NQO1, and Hoechst. Scale bar, 1 mm. B, Representative images of the olfactory nerve layer (ONL), glomerular layer (GL), and external plexiform layer (EPL) stained with GFAP, NQO1, and DAPI. White dashed circles indicate glomeruli. Scale bar, 200 μm. C,E, Areas of GFAP-positive cells in the ONL and GL. (n = 3, one-way ANOVA followed by Dunnett’s post hoc test. **p < 0.01, ***p < 0.001). D,F, Comparison of the normalized values of the density of GFAP-positive cells between NQO1-positive and NQO1-negative regions in the ONL and GL. (n = 3, Welch’s t-test).

**Figure 5-figure supplement 1.**
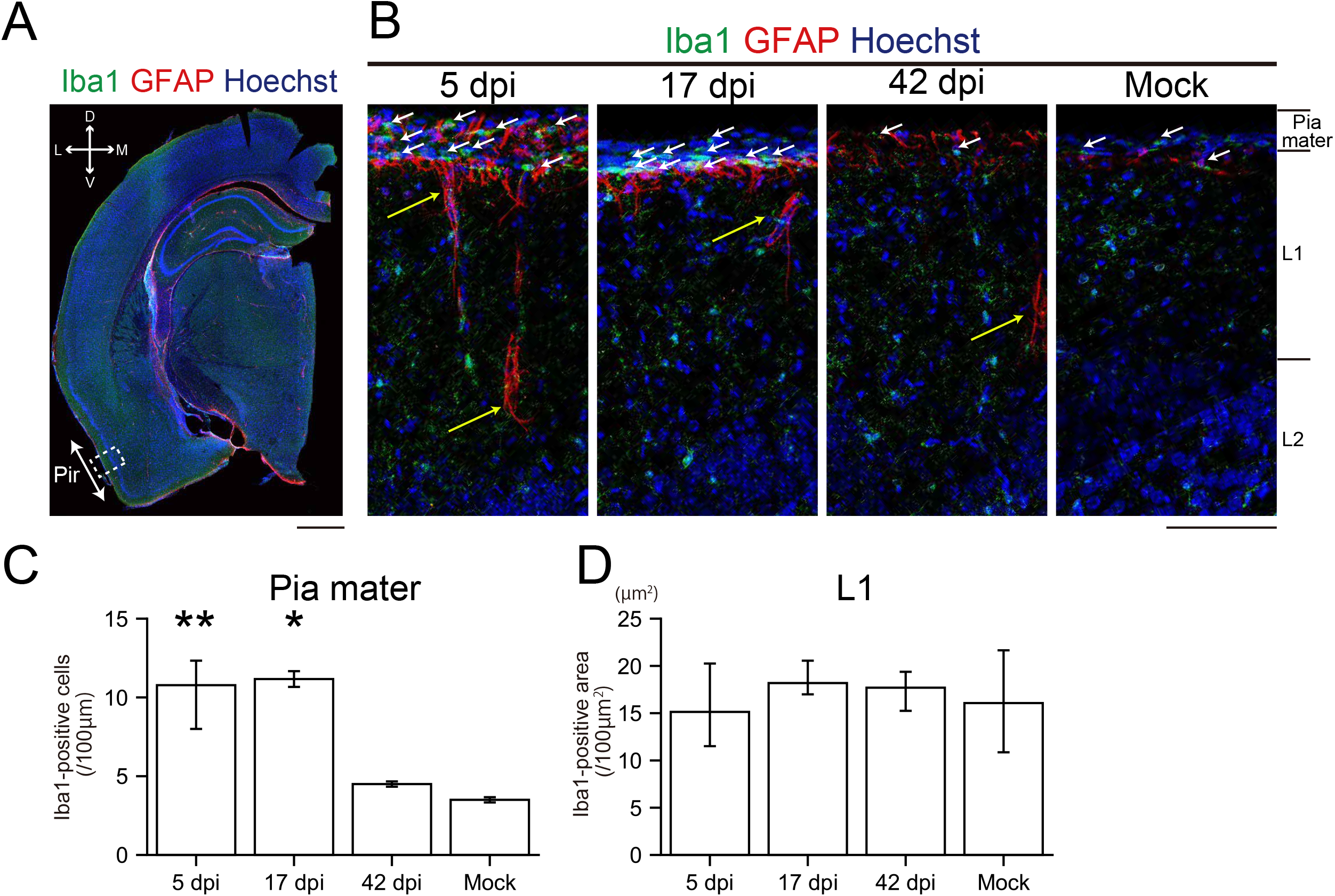
Glial activities in the piriform cortex (PC) due to SARS-CoV-2 infection. A, Representative coronal image of the cerebral hemisphere stained with ionized calcium binding adaptor molecule 1 (Iba1), GFAP, and Hoechst. The white arrow indicates the PC and the white dashed square is a region of interest in the study. Scale bar, 1 mm. B, Representative images of the layers of pia mater, layer 1 (L1), and layer 2 (L2) stained with Iba1, GFAP, and Hoechst at 5, 17, and 42 dpi and mock. White and yellow arrows show Iba1- and GFAP-positive cells, respectively. Scale bar, 100 μm. C, Quantitative analysis of Iba1-positive cells in the pia mater. (n = 3, one-way ANOVA followed by Dunnett’s post hoc test. *p < 0.05, **p < 0.01). D, Iba1-positive densities in L1. (n = 3, one-way ANOVA followed by Dunnett’s post hoc test).

